# Introducing a new methodology to improve the predictive value of ancient woodland indicator species for landscape history

**DOI:** 10.1101/2023.10.03.560441

**Authors:** Alistair Baxter, Julian Forbes-Laird

## Abstract

Certain species of vascular plants have been proposed as indicators of very long established woodland. In the England and Wales, woodland present continuously since at least 1600AD is known as ‘ancient woodland’, with the plants concerned being referred to as ‘ancient woodland indicator species’. A number of authors have considered this topic from differing angles, and the concept has been validated, though with problems identified. The authors introduce a new method, WISDOM, for recording and analysing indicator assemblages, and test this tool with a comparative assessment of their distribution and occurrence in a known ancient woodland and a known secondary woodland. This paper reports the results of this assessment, discussing also the evidential weight that can be attached to different species of indicator plants. A literature review is included, with the authors concluding that their findings are consistent with several strands of previous research.

## Introduction

The overall concept of *Ancient Woodland Indicator Species* (AWIS) is that certain woodland plants are slow colonists, therefore requiring very long-term woodland cover in order to establish (**Rose 1999**). Where such woodland predates 1600AD, and has been present continuously since that date, it is classed in England and Wales as Ancient Woodland, hence such plants are known as “Ancient Woodland Indicators”. Ancient Woodland in the UK generally has a high status within the planning system as *Irreplaceable Habitat*.

**Rose & O’Reilly 2006** compiles geographically relevant lists of the vascular plants considered by botanists to indicate the presence of very long-established woodland. According to Rose & O’Reilly, 185 plants are ancient woodland indicator species (AWIS). The authors of this paper add one additional species, Coralroot Bittercress *Cardamine bulbifera*.

Coralroot Bittercress has very strong fidelity to woodland and a very slow colonisation rate (it propagates by deadfall seed with low recruitment); these are paradigm attributes of the AWIS hypothesis. Whilst Coralroot Bittercress was omitted from the Rose & O’Reilly listing due to its rarity, in our view this merely reinforces its usefulness where found.

AWIS comprise both woody and non-woody species, with the latter subdividing further into grasses, sedges, rushes, ferns, and wildflowers.

There are several difficulties with the AWIS hypothesis.

1. It is known that the fidelity of many indicator species to woodland is at best questionable (e.g. Bitter-vetch *Lathyrus linifolius*).
2. Many other species have relatively rapid means of colonisation which enable them to occupy secondary woodland in decadal, rather than centennial timeframes (e.g. Bluebell *Hyacinthoides non-scripta*).
3. Some species are more strongly reflective of a prior alternative habitat to woodland (e.g. Bilberry *Vaccinium myrtillus*) and are, therefore, more correctly interpreted as contra-indicators.
4. The presence of a single or very low number of specimens of a particular species should generally attract limited significance, as they could be there as a result of chance: by definition, chance occurrence does not indicate the duration of woodland cover.
5. Many AWIS can lie dormant during long periods of unfavourable land conditions, including surviving below ground in various vegetative forms on the centennial scale. Bye Wood, at Winsford, Exmoor, provides a recent example. Land at this location had been an open hilltop for centuries prior to landscape restoration work triggering the emergence of a substantial bluebell carpet^1^. For this reason, AWIS cannot be used to indicate <u>continuity</u> of woodland.

### Differing perspectives on the AWIS hypothesis

In the UK, AWIS are used to assist in the identification of ancient woodland, including where woodlands become listed in the Ancient Woodland Inventory compiled and maintained by Natural England, and equivalent bodies in Northern Ireland, Scotland and Wales.

In its Inventory Handbook^2^, Natural England is at pains to avoid setting a threshold of significance for the presence of AWIS:

> *The indicator species tool is inexact but simple: the more species recorded on the site, the more likely that site is to be ancient (the presence of [indicator species] does not prove a wood is ancient and neither does their absence prove recentness). However, there is no linear relationship between the precise degree of likelihood and the number of species present that can be routinely applied at site-level. There is no threshold species count above or below which the status of a wood becomes a certainty. For this reason, [botanical] information should be used as a supporting part of the wider investigation*.

With this in mind, we provide here a review of the investigations and conclusions of others who have examined this topic before us.

**Dzwonko 1993** investigated the influence of ancient woodlands on the floristic composition of young woodland communities in the Carpathian foothills by examining four ca. 70-year-old deciduous woods adjacent to three ancient oak-hornbeam and oak-pine woodlands. The latter were found to have acted as a source of woodland species diaspores, and showed a significant relationship between the distance to ancient woodlands and species composition in recent woods on rich brown soils. On poor soils, dense growth of *Carex brizoides* strongly inhibited successful colonization by woodland specialist plants in this lower nutrient environment.

**Grashof-Bokdam & Geertsema 1998** investigated the colonization success of forest plant species in woodland habitats in the Netherlands, and confirmed the importance of dispersal strategies, age of habitat, distance to seed sources, and former land use (consistent with the findings of **Dupouey et al 2002**) on the occurrence of different species.

**Norden & Appelqvist 2001** argues that dispersal capacity is the key factor that defines the effectiveness of an indicator species. They propose that species with low dispersal capacity, such as terrestrial molluscs, some vascular forest plants (being the grouping that concerns us here), and certain bryophytes and lichenized fungi, may be the best indicators of long-term habitat persistence in forests. Species with known high dispersal factors should not be considered reliable indicators.

This approach is strongly consistent with that set out in **Thompson et al 2003**, which found that only around one third of suggested vascular plant species were reliable indicators of woodland habitat persistence. Whilst their study considered only one county in England, their conclusions are likely relevant to lowland Britain generally.

Meanwhile, **Rolstad et al 2002** set out a critique of the wider concept of indicator species, highlighting examples that demonstrate its limitations. Their study concludes by following **Lindenmayer et al 2000** in suggesting that structural indicators (i.e. the composition of forest stands) are a better guide to long-term habitat persistence than the presence or absence of species indicators. Clearly, this approach can only work in landscapes that lack a history of forest clear-felling and/ or of overplanting, and hence is of minimal, if any assistance in the British context.

Of greater relevance are the findings reported by **Petit et al 2004**, who analysed the richness of Ancient Woodland Indicator Species in British woodlands, testing the hypothesis that this is limited by woodland area, shape, and spatial isolation. They also investigated whether the lowland region would respond differently compared to the upland region.

Their results showed that indicator species richness in the lowlands was mainly influenced by woodland size and connectivity (e.g. the presence of connecting hedgerows). In contrast, indicator species richness in the uplands was related to habitat quality (Ellenberg scores), thereby echoing Dzwonko 1993, with no significant effect identified in relation to a woodland’s landscape context.

**Rotherham et al 2009** presents the results of a survey conducted to assess the use of ancient woodland indicator lists in the UK. This study found a general lack both of species thresholds and of weightings to determine whether a site is ancient woodland. While they found there are ca. 200 species listed on different indicator lists, few species are common to more than a quarter of them.

The methods used to develop indicator lists were found to be somewhat ad hoc, with some based on expert opinion, others on field surveys, and others on adaptation of existing lists. Few lists either use adequate, let alone extensive field surveys, or cross-referencing of field to archive botanical data. This study also raised concerns about the robustness of some existing lists, and the uncritical use of indicators in recent planning inquiries in England.

**Kelemen et al 2012** examined the effects of land-use history and current management practices on herbaceous plant composition in European woodlands, based on a study conducted in the Bakony Mountains of Western Hungary. The researchers compared the herbaceous plant assemblage in ancient forests, to that in both older and younger recent forests. They also looked for life-history traits that could explain the differences in floristic composition.

The results showed that plant species differed between all three categories of forest. The researchers classified the herbaceous species into four groups based on their ecological traits, and found that the group which included geophytes and herbs flowering in early spring were indicators of ancient forest stands. This is consistent with the generality of the AWIS hypothesis.

Recent, post-agricultural woodland was found to be largely dominated by competitive species, further supporting the AWIS hypothesis in broad terms.

Consistent with both Grashof-Bokdam & Geertsema 1998 and Dupouey et al 2002, this study highlights the importance of land-use history, as well as current management practices, in shaping the floristic assemblage of European woodlands.

**Kirby et al 2013** investigated the traits of ancient woodland indicator species to examine trait commonality, as well as how distinguishable any trait profile is from other woodland plants.

Their results show that AWIS are generally short, perennial plants with high seed weight. Rarer AWIS have a more distinct trait profile (poor dispersal ability and adaptation to low light availability during the growing season) than more common species. This supports our inclusion of Coralroot Bittercress as an indicator.

In strong agreement with Norden & Appelqvist 2001, Kirby et al identified poor dispersal ability as a key factor in confining AWIS to ancient woodlands. Phylogeny, however, was not found to be a factor in determining AWIS status, an interesting conclusion.

This study also found that the association between AWIS and ancient woodland habitat depends on landscape context, per Petit et al 2004 with regards to lowland Britain.

Overall, these findings provide a better understanding of why AWIS are found in ancient woodlands and can help in the identification and conservation of these habitats.

Finally, **Swallow et al 2020** used a statistical model to examine field work results, testing plant species diversity according to the alpha-beta-gamma diversity schema proposed by **Whitaker 1972**.

Their study examined three types of woodland: ancient semi-natural, ancient replanted, and secondary (recent) woodland. Their approach successfully distinguished between plant associates of the three woodland types (per Kelemen et al 2012).

### Proposed new methodology

Whilst the literature clearly finds merit in the use of plant indicators in historic landscape investigation, and as markers for biodiversity, it also identifies a number of problems and shortcomings. To seek to address these issues (at least in part), we propose here a new assessment method in the form of WISDOM (for *Woodland Indicators: Strength, Distribution and Occurrence Matrix*).

WISDOM is a list of the 185+1 indicators (see Table 3, following), which in spreadsheet form is selectable on a county or regional basis. Woodlands can be assessed using the tool either as a single compartment, or broken down into constituent parts.

It is known that some AWIS are more reliable than others as indicators of very long-established woodland. Four examples illustrate the point:

- Herb Paris *Paris quadrifolia* has very high fidelity to woodland and is a very slow colonist. Its presence is a strong indicator for very long-established woodland.
- Oak Fern *Gymnocarpus dryopteris* has high fidelity to woodland and is readily destroyed by grazing. It is a slow-growing plant and so cannot re-establish easily once lost from a site. It is also a strong indicator.
- Bluebell *Hyacinthoides non-scripta* is well-known for colonising even relatively young woodland. It is, therefore, a weak indicator.
- Great Burnet-saxifrage *Pimpinella major* is a plant of several different habitats, of which woodland is only one, and disperses well by freely-set seed: it is also a weak indicator.

This evidential variability of differing indicator species confirms that assessment of indicator presence should be nuanced.

The WISDOM methodology assigns to each of the 186 indicator species an evidential weighting, *Strong, Moderate* or *Weak*. We summarise the three strength categories as follows:

- Strong Woodland specialist & slow coloniser due to dispersal method
- Moderate Woodland specialist but more rapid coloniser, *or* Slower coloniser of more than one habitat type
- Weak Generalist and/ or swift to colonise

The strength assignment has been derived by a three-step process:

1. We assigned S-M-W to each plant based on occurrences within the British *Southern Ancient Woodland Inventory*, following a review of the updated local inventory reports of which that is comprised. Species with typically a reduced frequency of occurrence were given a stronger weighting, and those with a greater or high frequency of occurrence were given a weaker weighting.
2. We reviewed Rose & O’Reilly 2006 for all indicators which we had initially classified as *Strong* to check their fidelity to woodland. Where an indicator was noted as occurring in one or more alternative habitats, it was downgraded to *Moderate*.
3. Finally, we revisited the list of *Moderate* strength indicators, undertaking a further adjustment: a) if habitat is confined or largely confined to woodland, and where colonisation rate is slow, the plant was elevated to *Strong*; and b) if a plant’s habitats are varied and/ or where its colonisation rate is moderate or swift, the plant was downgraded to *Weak*.

The final classification of plants as S-M-W rests on a combination of reported occurrences in ancient woodland; on habitat preferences; and on colonisation strategy and success. Whilst we recognise that species assignments will always be open to some debate, the chosen categories provide a reasoned starting position to enable testing of the tool by other practitioners. The authors would welcome comments on this topic to their correspondence address.

The area count of *Strong, Moderate* and *Weak* WISDOM scores are reported in the format *x*.*y*.*z*, thereby deriving not only the total number of indicator species present (alpha diversity, after Whittaker), but also their relative strength as signposts towards woodland antiquity.

### Interpretation of WISDOM scores

The *Strong-Moderate-Weak* classification of all 186 plants within WISDOM is 66.85.35. However, because AWIS have a significant element of local distinctiveness, no district, county, or regional AWI list contains all 186 species. By way of example, the Sussex AWI list contains 100 plants, with a WISDOM score of 38.51.11.

In general terms, species aggregation is a function of time, area and land management. Where both types of woodland are of the same age, a small, semi-natural woodland may well contain greater AWI variety than a larger woodland under conifer plantation.

However, factors such as an excessive deer population, or recent, heavily mechanised clear-felling, are also relevant, as is whether a conifer plantation has been thinned, as opposed to forming a dense, closed canopy.

Relative abundance is, therefore, an important consideration when assessing the significance of indicators. The WISDOM spreadsheet includes a D-A-F-O-R scale assessment (Dominant, Abundant, Frequent, Occasional, Rare). The DAFOR scale can be applied at whole wood or compartment level. The use of occurrence recording permits a further layer of interpretation: in a large woodland, the presence of an indicator in a single location, may suggest a chance occurrence of, therefore, low evidential weight (even if the indicator weighting is Strong).

The authors have resisted the temptation to set hard thresholds for WISDOM scores, as it is important to interpret botanical survey results within the context of the case-specific woodland to which they relate, as well as in light of other evidence streams (the site’s cartographic record, for example).

However, based on our experience of undertaking numerous woodland botanical surveys using WISDOM, we attach a degree of significance to the presence of 3 or more *Strong* indicators, and higher significance still where there are five or more present (in both these cases, *Moderate* and *Weak* indicators are likely present too). We consider that a WISDOM score of around 5.10.5 reflects a significant assemblage, albeit one to be treated with due caution for the reasons given already.

### Comparative assessment study

In order to directly test both the WISDOM tool, and also the AWIS hypothesis under British conditions, in 2022 the authors undertook a research project comprising a comparative floristic assessment of two woodlands. Both woodlands were visited during the prime recording season for AWIS (April to June inclusive).

The subject woodlands were identified based on cartographic research, enabling their landscape history to be determined with a high level of accuracy. The details of the two study woodlands are set out in Table 1.

**Table 1.**
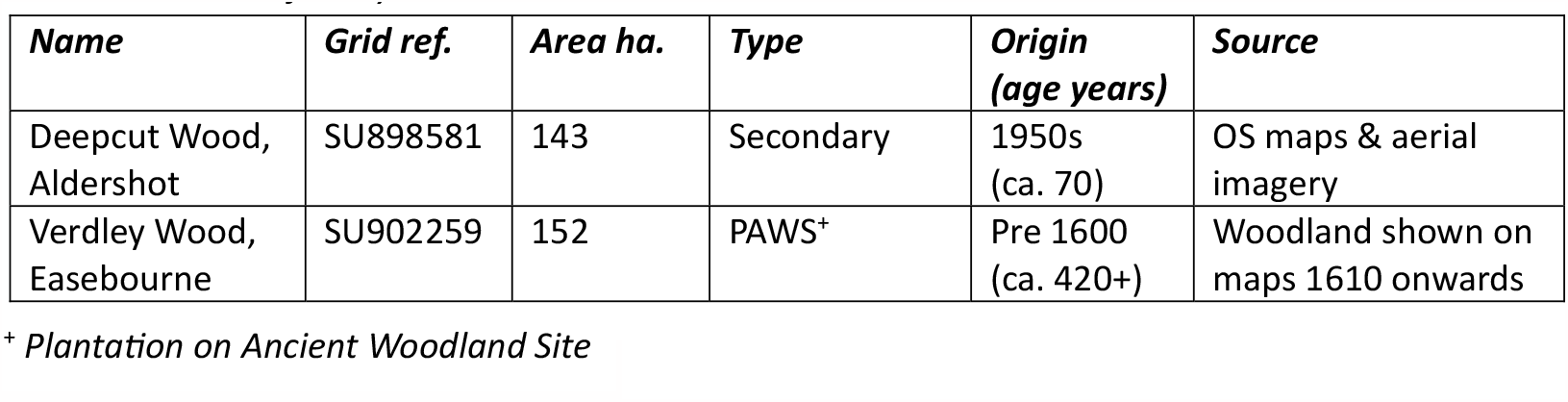
Details of study woodlands. ^+^ Plantation on Ancient Woodland Site

#### Deepcut Wood

Deepcut Wood (Figure 1) lies at the eastern edge of Frimley and southeast of Camberley. It is known to have been under heath until the 1950s, before it was planted for forestry; it is, therefore, Secondary Woodland).

**Figure 1.**
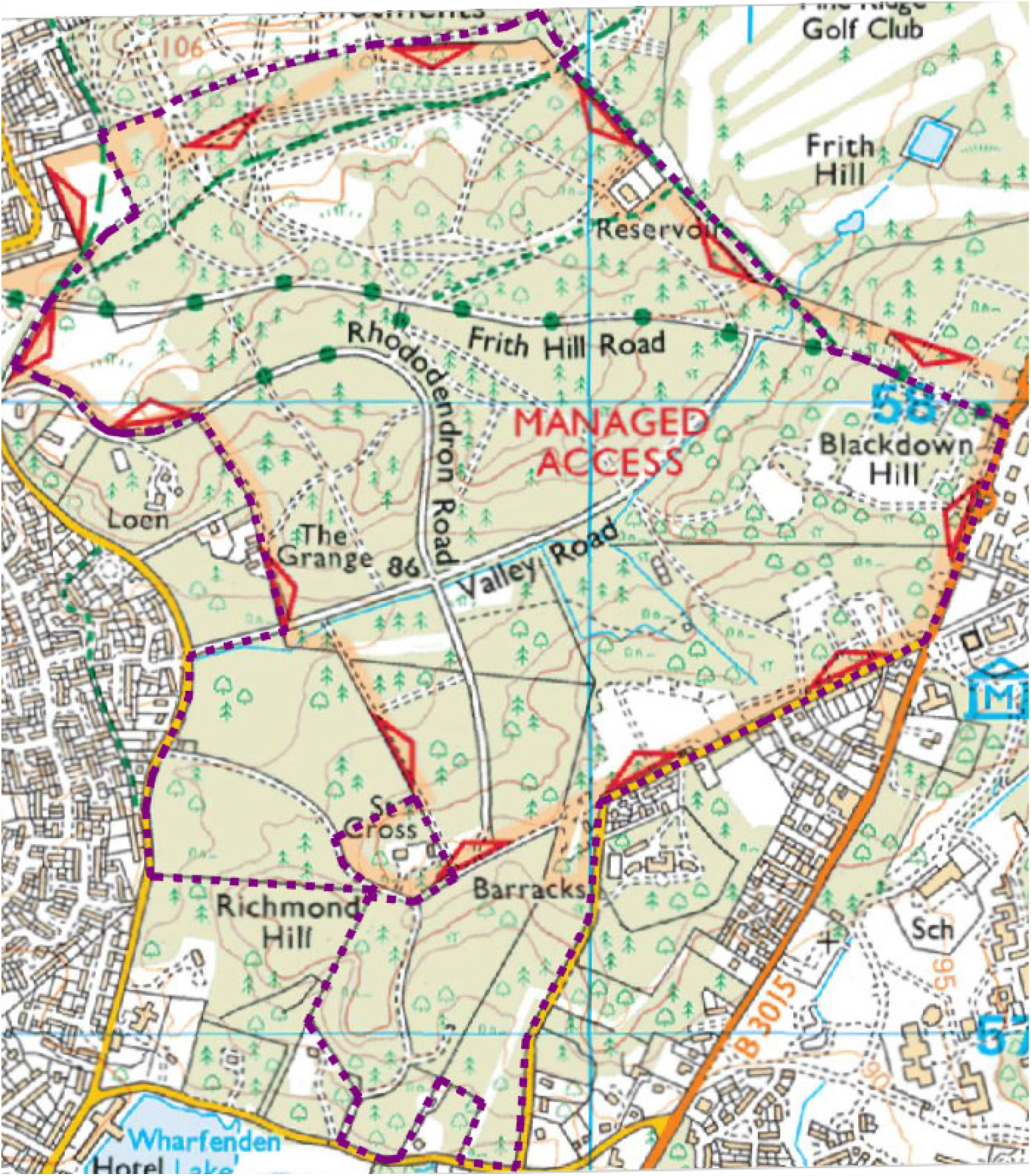
Ordnance Survey extract showing Deepcut Wood; Sylvan survey area in dashed outline.

Deepcut Wood is used extensively for military training, with parts of it being frequently closed to public access. It is, however, generally open to the public, and provides a valued leisure resource to the local community, including dog-walkers.

Deepcut Wood includes three named hills, Frith Wood, Blackdown and Richmond, but none are more than minor variations in the local topography. To the north, the edge of the wood achieves an elevation of 105m, falling to 86m centrally, before rising to 95m in the south. The low point sits on a roughly west to east ride called Valley Road.

The wood stands on free-draining sandy soil. There is only one watercourse present, which appears to be man-made.

According to the National Forest Inventory (Figure 2), Deepcut Wood is majority conifer, though with substantial areas of priority habitat broadleaved woodland is also present. Indeed, our survey found more broadleaf present than indicated by the NFI. Very little evidence of sylviculture was found during our survey.

**Figure 2.**
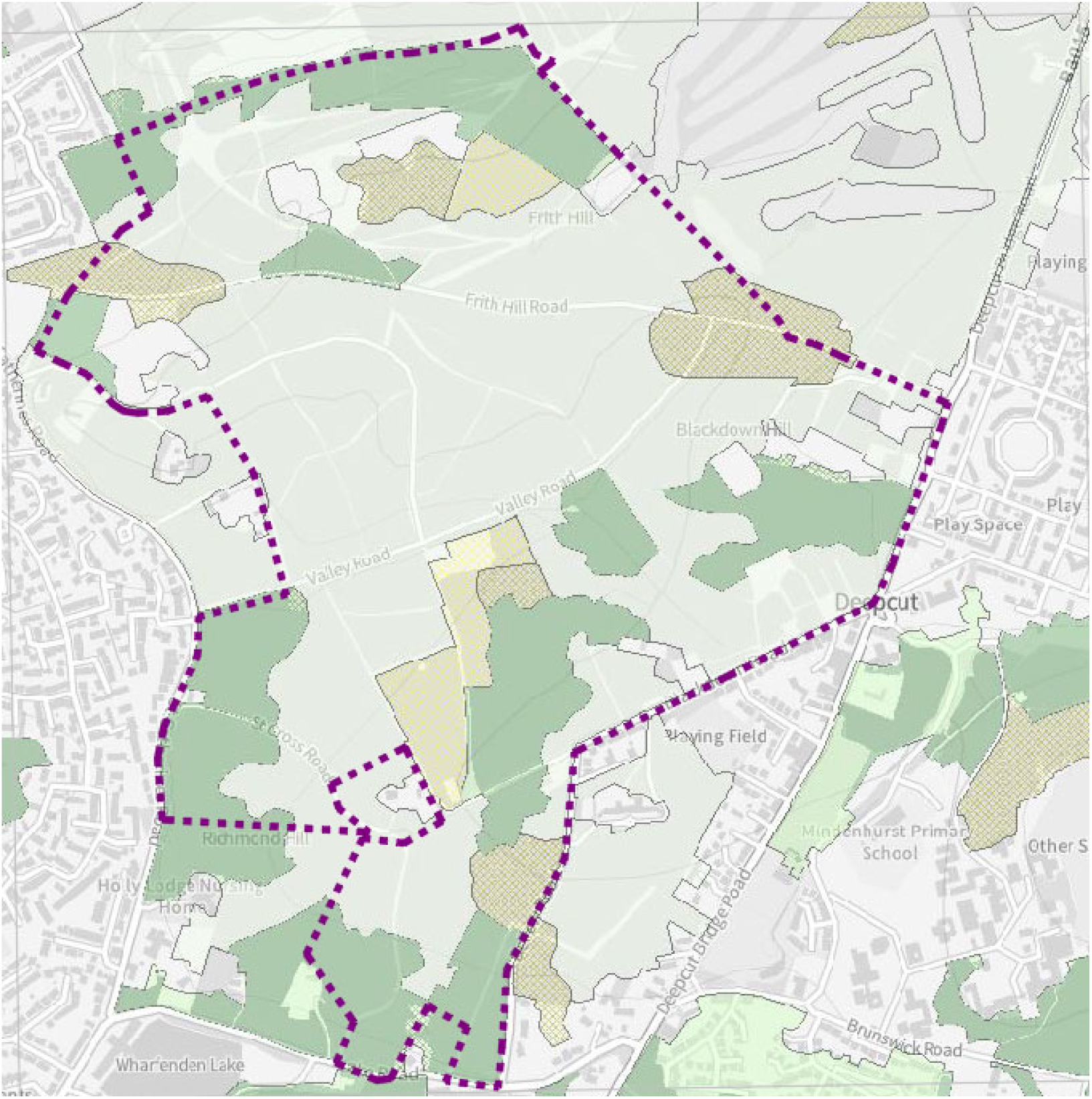
Extract from MAGIC^3^ showing National Forest Inventory data; light green indicates conifer, dark green indicates broadleaf.

There is minimal Ancient Woodland within the locality, though a small area of it lies ca. 110m to the west of Deepcut Wood. However, this is substantially separated from Deepcut Wood by housing, and accordingly, there is no significant colonisation pathway between the two woodlands for AWIS.

#### Verdley Wood

Verdley Wood (Figure 3) lies within the “Western Ridges” landscape character area (LCA) of West Sussex^4^. This LCA features an undulating dip-slope of typically wooded ridges, which rise steadily to the north of the Rother Valley before dropping rapidly away to a steep, deeply indented escarpment. The escarpment is incised by deep, occasionally ravine-like stream valleys.

**Figure 3.**
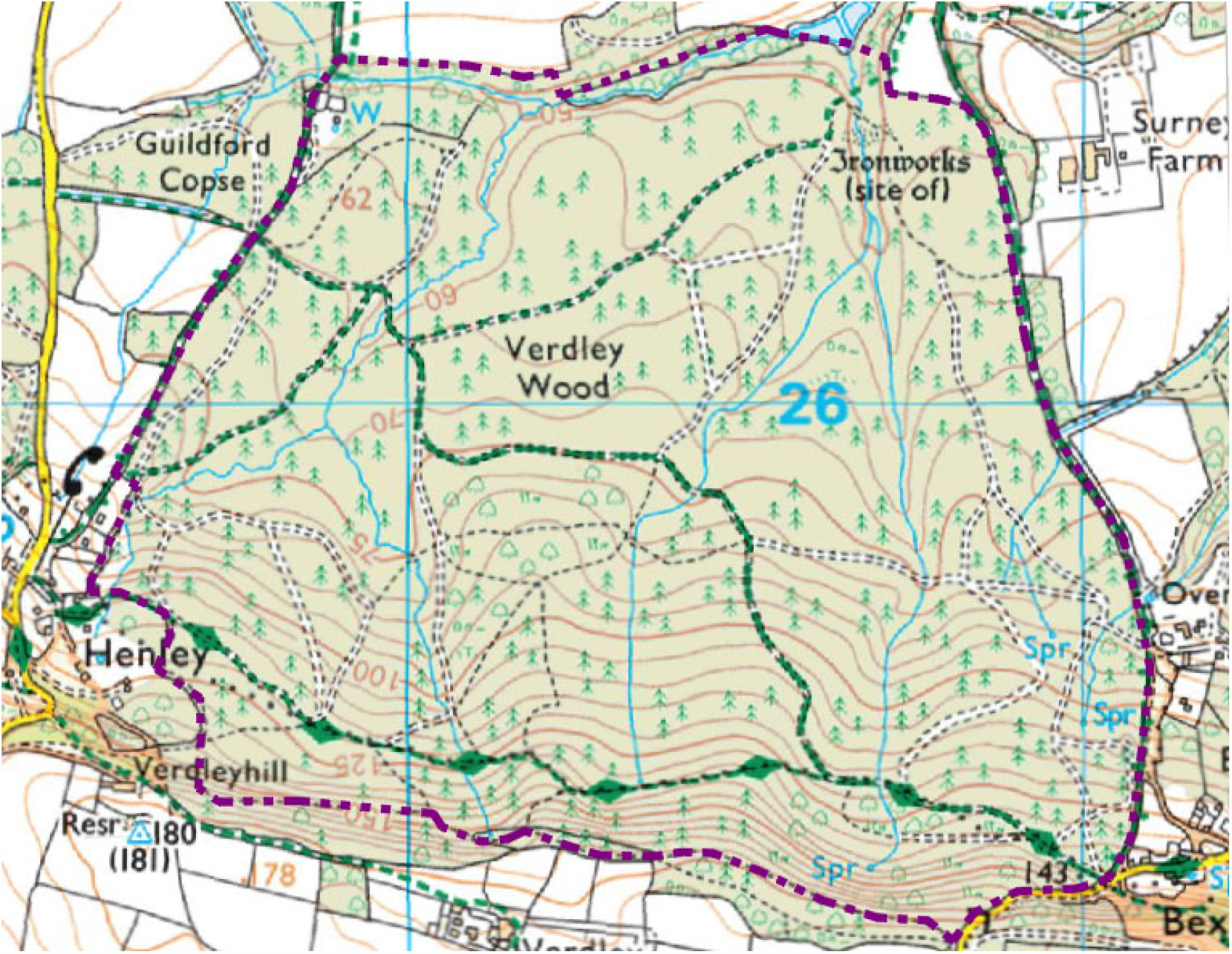
Ordnance Survey extract showing Verdley Wood; Sylvan survey area in dashed outline.

Verdley Wood lies substantially on the northerly, dip slope of the ridgeline. The wood is underlain by acid soils, including combination superficial deposits of clay/silt/sand as well as sand in isolation, and bedrock deposits of mudstone and sandstone.

The site is drained north to south by five watercourses, which are incised in places, with three of the five rising as identifiable natural springs. The five watercourses feed into a larger watercourse at the foot of the dip, which flows west to east roughly along the wood’s northerly boundary.

Cartographic research undertaken by the authors has identified Verdley Wood on maps dating to 1610AD (Figure 4), which supports Natural England’s decision to include it within the Ancient Woodland Inventory. However, it is included within the Inventory as a *Plantation on Ancient Woodland Site* (PAWS), with the National Forest Inventory (Figure 5) identifying it as almost wholly under coniferous planting.

**Figure 4.**
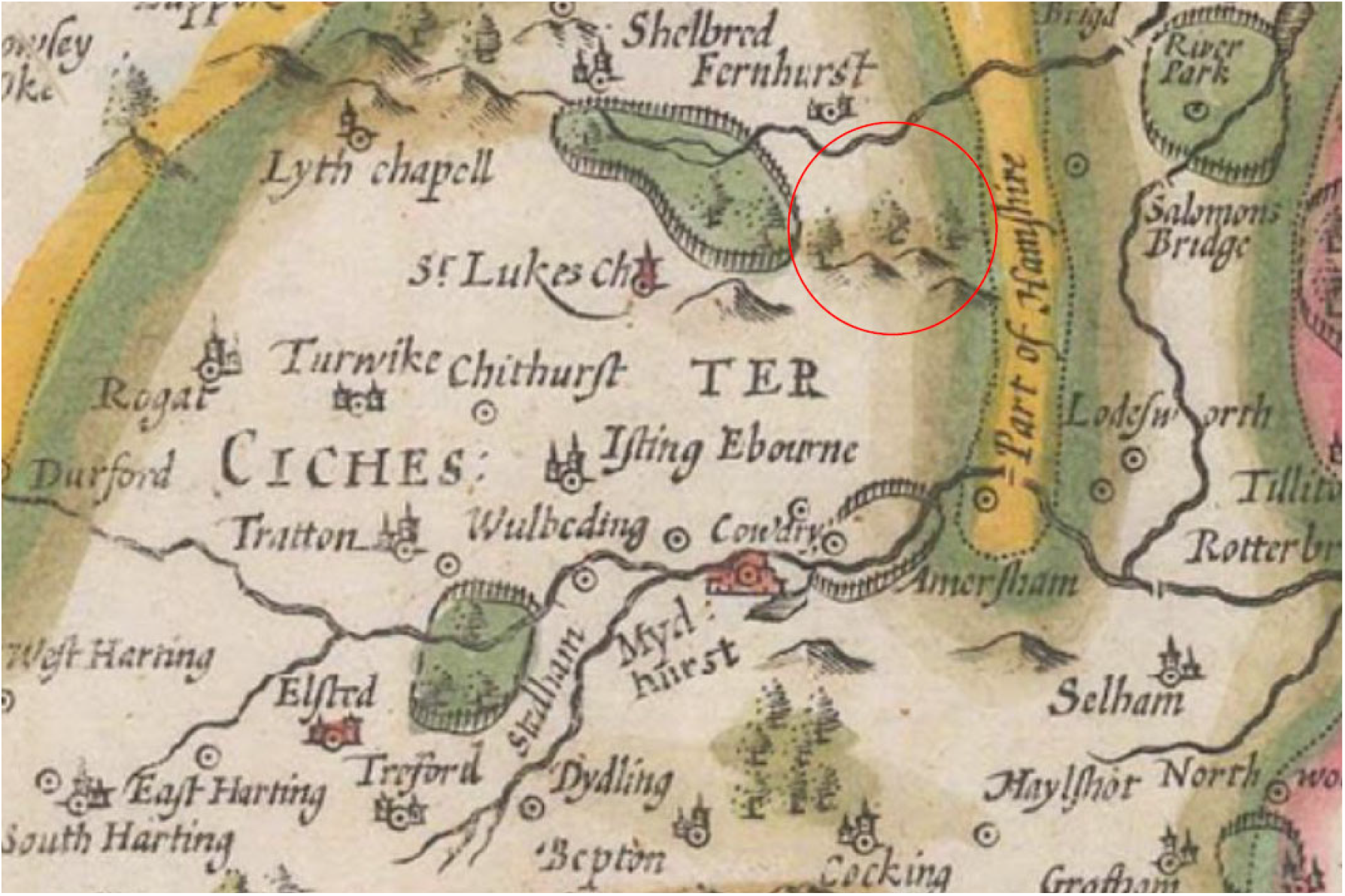
Extract from Speed’s map of Sussex, 1610; site locus circled (note the presence of forest)

**Figure 5.**
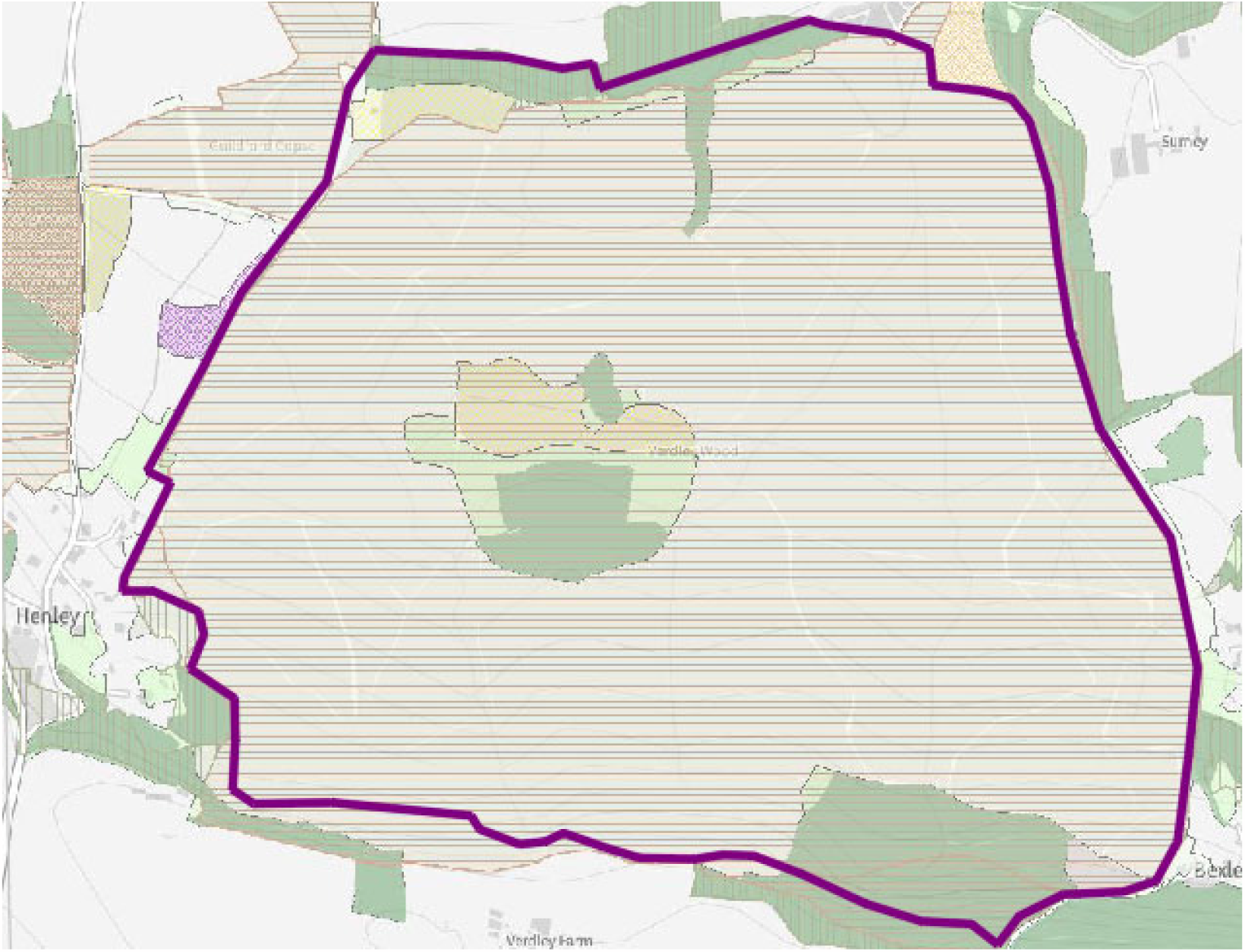
Extract from MAGIC showing National Forest Inventory data; fawn conifer, dark green broadleaf; hatching PAWS.

The conversion to conifer appears to have happened after 1910: at that time, the Ordnance Survey Six Inch map (Figure 6) records the presence of broadleaved woodland. The 1:25K OS map of 1958 (Figure 7) shows mixed conifer and broadleaf.

**Figure 6.**
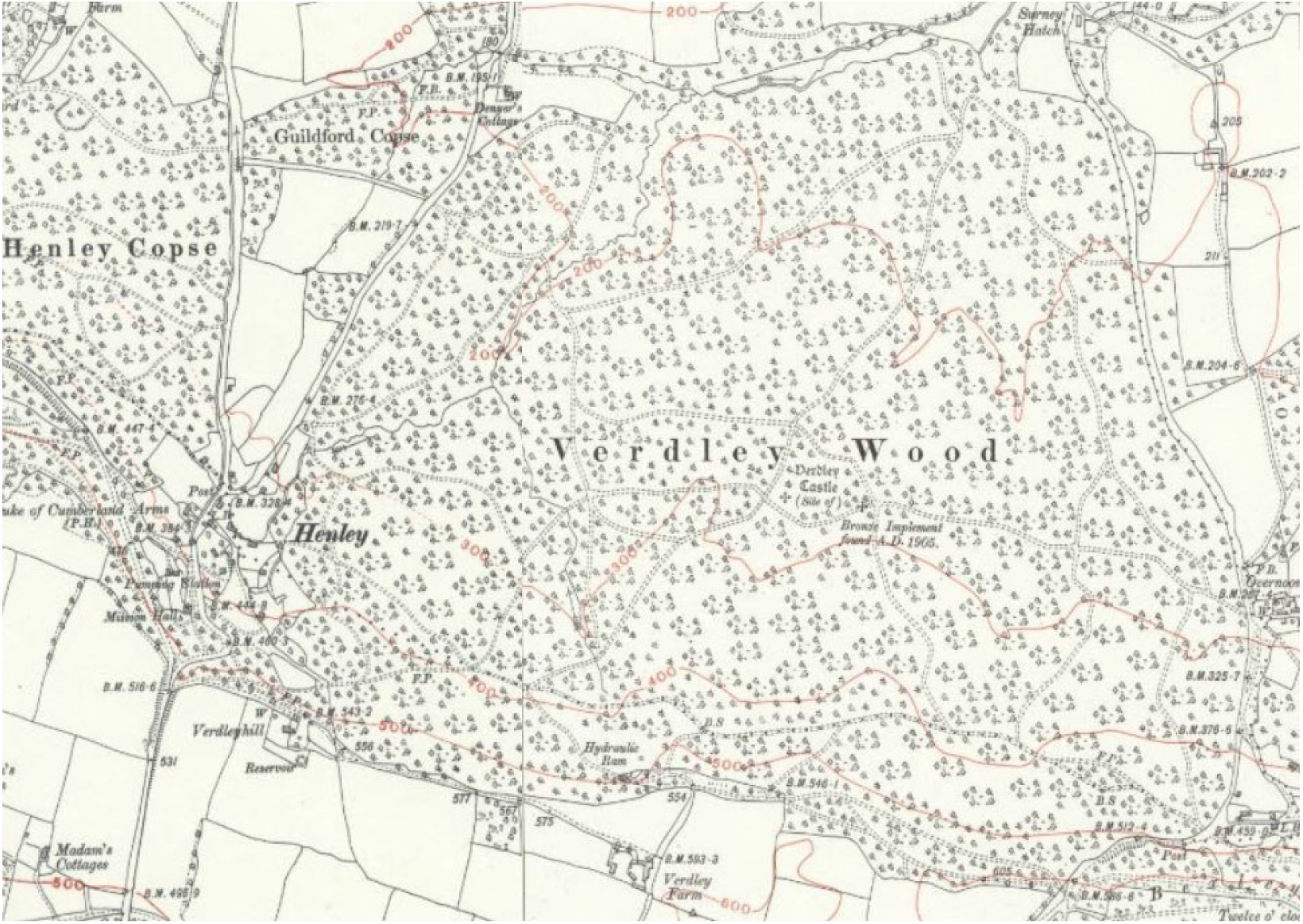
Extract from Ordnance Survey 6” map 1910 showing broadleaf tree cover.

**Figure 7.**
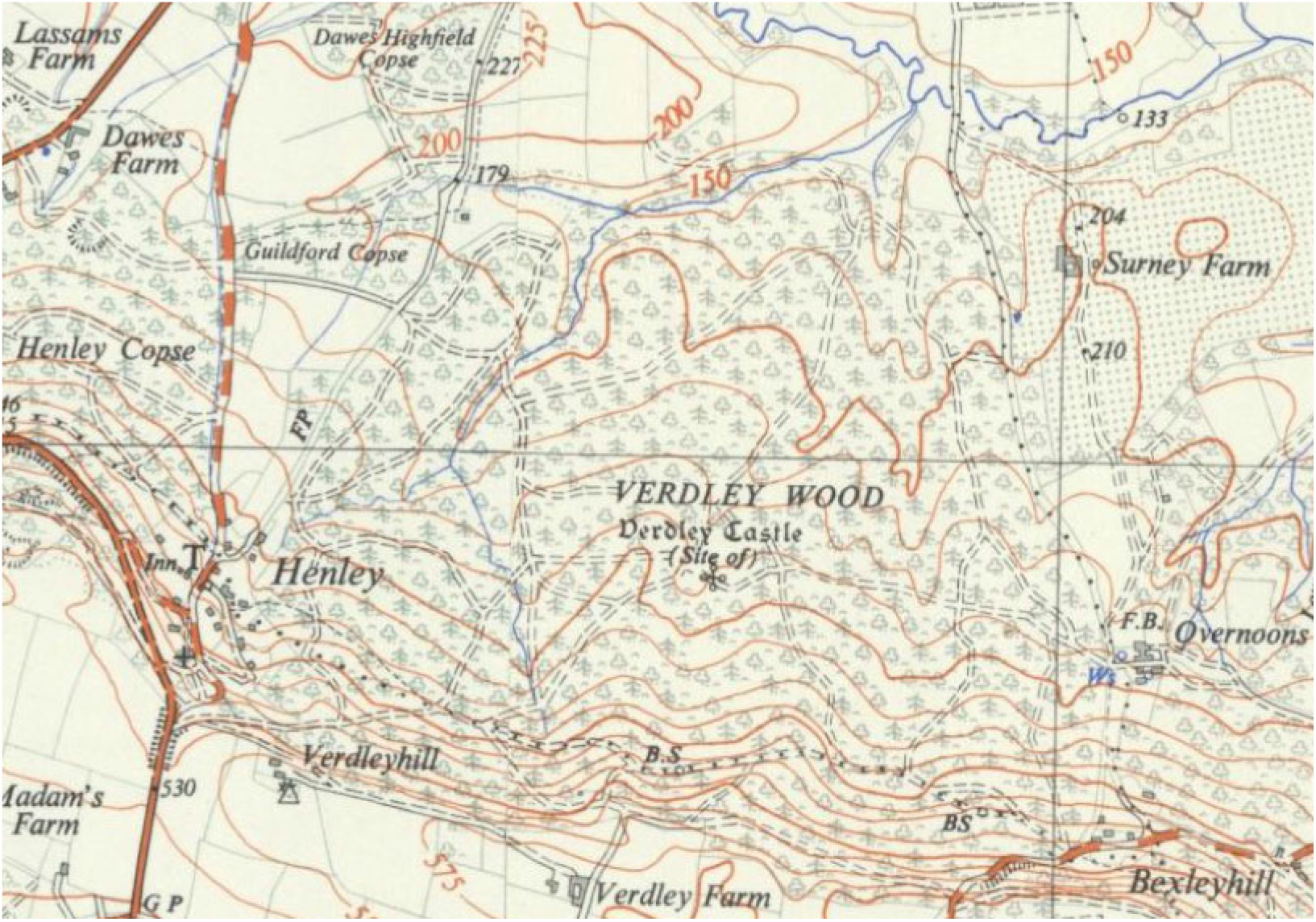
Extract from Ordnance Survey 1-25K map 1958 showing mixed tree cover.

Active management was found to be in hand, comprising forestry in the conifer plantation areas, and sylviculture in the minority broadleaved areas. Public access appears to be relatively infrequent, presumably due to its location somewhat away from substantial population centres.

Ancient Woodland is strongly prevalent in the local area, including lying contiguous and near contiguous to Verdley Wood in several places. This landscape-scale mosaic of Ancient Woodland provides durable pathways for inter-woodland species exchange, including AWIS.

### Assessment methodology

The two study woodlands were visited in April and June 2022, and were subject to a Level 2 Walkover Survey (**Kirby & Hall 2019**). Five days were spent in Verdley Wood, and four in Deepcut Wood. Rides, substantial paths, and watercourses (collectively here termed “walk-routes”) were used as proxy transects, due to the known function of edge and riparian habitats as hotspots for floristic diversity. In a few cases, transects were made across larger compartments where walk-routes were absent.

The distance covered within each of the woodlands was similar (Figure 8). Approximately 21km were walked in Verdley Wood, of which ca. 4.5km was along watercourses. The figures for the slightly smaller Deepcut Wood are 19km and 850m respectively. The shorter watercourse distance in Deepcut Wood reflects its general lack of these features.

**Figure 8.**
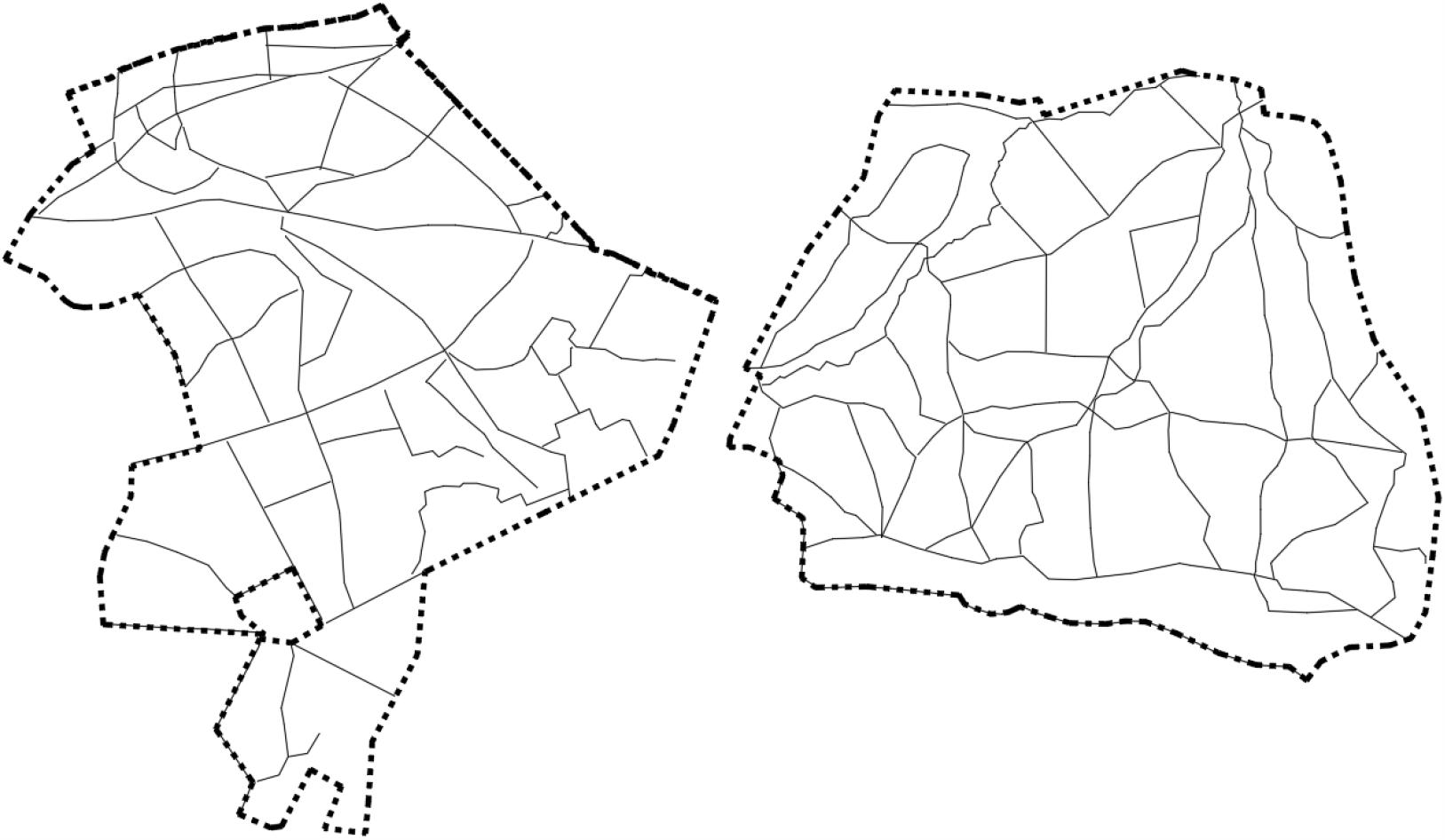
Ground plan showing Sylvan survey area & walk-routes; Deepcut left and Verdley right.

AWIS were recorded using WISDOM, with occurrence of each species assessed with the DAFOR scale.

As noted, Deepcut Wood is located in Surrey, whilst Verdley Wood lies within Sussex; the appropriate county indicator list (Rose & O’Reilly 2006) has been used for each. Both the Surrey and Sussex AWIS lists contain 100 species, such that the total species count for each woodland is also the percentage of AWIS found relative to the respective county list.

### Summary of Findings

Our findings for each of the study woodlands are set out in Table 4 (Deepcut Wood) and 5 (Verdley Wood), following, but are summarized in Table 2.

**Table 2.** Summary of findings for each study woodland.

| **Name** | **Strong** | **Moderate** | **Weak** | **WISDOM**<br>**score** | **Total** | **County %** |
| --- | --- | --- | --- | --- | --- | --- |
| Deepcut Wood | 3 | 11 | 5 | 3.11.5 | 19 | 19 |
| Verdley Wood | 11 | 28 | 4 | 11.28.4 | 43 | 43 |

**TABLE 3.** ANCIENT WOODLAND INDICATORS + SYLVAN WEIGHTING. Species list after Rose & O’Reilly 2006

| Common Name | Scientific Name | Strong | Moderate | Weak |
| --- | --- | --- | --- | --- |
|  |  | Woodland specialist +<br>slow colonizer | Woodland specialist<br>but known to colonize<br>secondary woodland | Generalist and/ or<br>swift colonizer |
|  | Count | 66 | 85 | 35 |
| * Caution required if specimen found within woodland edge: risk of garden escape or seed transport from outside woodland |  |  |  |  |
| Alder Buckthorn | *Frangula alnus* |  | x |  |
| Allseed | *Radiola linoides* |  | x |  |
| Alternate-leaved Golden-saxifrage | *Chrysosplenium alternifolium* | x |  |  |
| Angular Solomon's-seal | *Polygonatum odoratum* | x |  |  |
| Aspen | *Populus tremula\** |  | x |  |
| Barren Strawberry | *Potentilla sterilis* |  | x |  |
| Bastard Balm | *Melittis melissophyllum* | x |  |  |
| Bearded Couch | *Elymus caninus* |  | x |  |
| Beech Fern | *Phegopteris connectilis* | x |  |  |
| Betony | *Stachys officinalis* |  | x |  |
| Bilberry | *Vaccinium myrtillus* |  | x |  |
| Bird Cherry | *Prunus padus* |  |  | x |
| Bird's-nest Orchid | *Neottia nidus-avis* | x |  |  |
| Bitter-vetch | *Lathyrus linifolius* |  |  | x |
| Black Bryony | *Tamus communis* |  | x |  |
| Black Currant | *Ribes nigrum\** |  | x |  |
| Bloody Crane's-bill | *Geranium sanguineum* |  | x |  |
| Bluebell | *Hyacinthoides non-scripta* |  |  | x |
| Broad-leaved Helleborine | *Epipactis helleborine* | x |  |  |
| Buckthorn | *Rhamnus catharticus* |  | x |  |
| Bush Vetch | *Vicia sepium* |  | x |  |
| Butcher's Broom | *Ruscus aculeatus* | x |  |  |
| Chaffweed | *Anagallis minima* |  | x |  |
| Climbing Corydalis | *Ceratocapnos claviculata* |  | x |  |
| Columbine | *Aquilegia vulgaris\** | x |  |  |
| Common Adder's-tongue | *Ophioglossum vulgatum* |  | x |  |
| Common Cow-wheat | *Melampyrum pratense* |  | x |  |
| Common Dog-violet | *Viola rivinana* |  |  | x |
| Common Figwort | *Scrophularia nodosa* |  |  | x |
| Common Twayblade | *Listera ovata* |  | x |  |
| Common Valerian | *Valeriana officinalis* |  | x |  |
| Common Wintergreen | *Pyrola minor* | x |  |  |
| Coralroot Bittercress | *Cardamine bulbifera* | x |  |  |
| Cornish Moneywort | *Sibthorpia europaea* |  | x |  |
| Crab Apple | *Malus sylvestris\** |  | x |  |
| Creeping Lady's-tresses | *Goodyera repens* | x |  |  |
| Creeping Soft-grass | *Holcus mollis* |  | x |  |
| Crested Cow-wheat | *Melampyrum cristatum* | x |  |  |
| Cuckoo Pint | *Arum maculatum* |  | x |  |
| Dewberry | *Rubus caesius* |  |  | x |
| Dog's Mercury | *Mercurialis perennis* |  |  | x |
| Dogwood | *Cornus sanguinea* |  |  | x |
| Downy Currant | *Ribes spicatum* |  | x |  |
| Eared Willow | *Salix aurita* |  |  | x |
| Early Dog-violet | *Viola reichenbachiana* | x |  |  |
| Early Purple Orchid | *Orchis mascula* | x |  |  |
| Elongated Sedge | *Carex elongata* | x |  |  |
| False Brome | *Brachypodium sylvaticum* |  | x |  |
|  |  | Woodland specialist +<br>slow colonizer | Woodland specialist<br>but known to colonize<br>secondary woodland | Generalist and/ or<br>swift colonizer |
| Field Maple | *Acer campestre* * |  |  | x |
| Field-rose | *Rosa arvensis* |  | x |  |
| Fingered Sedge | *Carex digitata* | x |  |  |
| Fly Orchid | *Ophrys insectifera* |  | x |  |
| Giant Bellflower | *Campanula latifolia* |  | x |  |
| Globeflower | *Trollius europaeus* |  | x |  |
| Golden-rod | *Solidago virgaurea* |  |  | x |
| Goldilocks Buttercup | *Ranunculus auricomus* | x |  |  |
| Great Burnet-saxifrage | *Pimpinella major* |  |  | x |
| Great Fescue | *Festuca gigantea* |  | x |  |
| Great Horsetail | *Equisetum telmateia* |  |  | x |
| Great Wood-rush | *Luzula sylvatica* |  | x |  |
| Greater Butterfly-orchid | *Platanthera chlorantha* | x |  |  |
| Greater Chickweed | *Stellaria neglecta* |  | x |  |
| Greater Stitchwort | *Stellaria holostea* |  | x |  |
| Greater Tussock Sedge | *Carex paniculata* |  |  | x |
| Green Hellebore | *Helleborus viridis* | x |  |  |
| Green-flowered Helleborine | *Epipactis phyllanthes* | x |  |  |
| Guelder Rose | *Viburnum opulus* * |  | x |  |
| Hairy Lady's Mantle | *Alchemilla filicaulis* |  | x |  |
| Hairy St John's-wort | *Hypericum hirsutum* |  | x |  |
| Hairy Wood-rush | *Luzula pilosa* |  | x |  |
| Hairy-brome | *Bromopsis ramosa* |  | x |  |
| Hard Fern | *Blechnum spicant* |  | x |  |
| Hard Shield-fern | *Polystichum aculearum* | x |  |  |
| Hart's-tongue | *Phyllitis scolopendrium* * |  | x |  |
| Hay-Scented Buckler-fern | *Dryopteris aemula* | x |  |  |
| Hazel | *Corylus avellana* |  |  | x |
| Heath Cudweed | *Gnaphalium sylvaticum* |  |  | x |
| Herb-Paris | *Paris quadrifolia* | x |  |  |
| Holly | *Ilex aquifolium* |  |  | x |
| Honeysuckle | *Lonicera periclymenum* |  |  | x |
| Hornbeam | *Carpinus betulus* * |  | x |  |
| Ivy Broomrape | *Orobanche hederæ* |  | x |  |
| Ivy-leaved Bellflower | *Wahlenbergia hederacea* |  |  | x |
| Juniper | *Juniperus communis* |  | x |  |
| Lady Fern | *Athyrium filix-femina* |  | x |  |
| Lady Orchid | *Orchis purpurea* |  | x |  |
| Large Bitter-cress | *Cardamine amara* | x |  |  |
| Large-leaved Lime | *Tilia playphyllos* |  |  | x |
| Lemon-scented Fern | *Oreopteris limbosperma* |  | x |  |
| Lesser Hairy Brome | *Bromopsis benekenii* | x |  |  |
| Lesser Pond Sedge | *Carex acutiformis* |  |  | x |
| Lesser Skullcap | *Scutellaria minor* |  | x |  |
| Lily-of-the-Valley | *Convallaria majalis* * | x |  |  |
| Marsh Violet | *Viola palustris* |  |  | x |
| May Lily | *Maianthemum bifolium* | x |  |  |
| Meadow Saffron | *Colchicum autumnale* | x |  |  |
| Melancholy Thistle | *Cirsium heterophyllum* |  |  | x |
| Mezereon | *Daphne mezereum* | x |  |  |
| Midland Hawthorn | *Crataegus laevigata* |  | x |  |
| Monk's-hood | *Aconitum napellus* | x |  |  |
| Moschatel | *Adoxa moschatellina* | x |  |  |
| Mountain or Nodding Melick | *Melica nutans* |  | x |  |
|  |  | Woodland specialist +<br>slow colonizer | Woodland specialist<br>but known to colonize<br>secondary woodland | Generalist and/ or<br>swift colonizer |
| Narrow Buckler-fern | *Dryopteris carthusiana* |  | x |  |
| Narrow-leaved Bitter-cress | *Cardamine impatiens* |  | x |  |
| Narrow-leaved Everlasting-pea | *Lathyrus sylvestris* |  | x |  |
| Narrow-leaved Helleborine | *Cephalanthera longifolia* | x |  |  |
| Narrow-leaved Lungwort | *Pulmonaria longifolia* | x |  |  |
| Narrow-lipped Helleborine | *Epipactis muelleri* | x |  |  |
| Nettle-leaved Bellflower | *Campanula trachelium* | x |  |  |
| Oak Fern | *Gymnocarpium dryopteris* | x |  |  |
| Opposite-leaved Golden-saxifrage | *Chrysosplenium oppositifolium* | x |  |  |
| Orpine | *Sedum telephium* | x |  |  |
| Oxlip | *Primula elatior* | x |  |  |
| Pale Sedge | *Carex pallescens* |  | x |  |
| Pear | *Pyrus communis* |  |  | x |
| Pendulous Sedge | *Carex pendula* * |  | x |  |
| Pignut | *Conopodium majus* |  | x |  |
| Polypody (all species) | *Polypodium vulgare (sensu lato)* | x |  |  |
| Primrose* | *Primula vulgaris* |  | x |  |
| Purple Small-reed | *Calamagrostis canescens* |  |  | x |
| Ramsons | *Allium ursinum* | x |  |  |
| Red Currant | *Ribes rubrum* * |  | x |  |
| Remote Sedge | *Carex remota* |  | x |  |
| Rowan | *Sorbus aucuparia* |  |  | x |
| Sanicle | *Sanicula europaea* |  | x |  |
| Saw-wort | *Serratula tinctoria* | x |  |  |
| Scaly Male-fern | *Dryopteris affinis* |  | x |  |
| Sessile Oak | *Quercus petraea* * |  | x |  |
| Slender St John's-wort | *Hypericum pulchrum* |  | x |  |
| Small Cow-wheat | *Melampyrum sylvaticum* | x |  |  |
| Small Teasel | *Dipsacus pilosus* | x |  |  |
| Small-leaved Lime | *Tilia cordata* * | x |  |  |
| Smooth-stalked Sedge | *Carex laevigata* | x |  |  |
| Snowdrop | *Galanthus nivalis* |  |  | x |
| Soft Shield-fern | *Polystichum setiferum* |  | x |  |
| Soft-leaved Sedge | *Carex montana* |  |  | x |
| Solomon's-seal | *Polygonatum multiflorum* | x |  |  |
| Southern Wood-rush | *Luzula forsteri* | x |  |  |
| Spindle | *Euonymus europaeus* |  |  | x |
| Spreading Bellflower | *Campanula patula* | x |  |  |
| Spurge-laurel | *Daphne laureola* | x |  |  |
| Square-stalked St John's-wort | *Hypericum tetraperum* |  | x |  |
| Stinking Hellebore | *Helleborus foetidus* |  | x |  |
| Stinking Iris | *Iris foetidissima* |  | x |  |
| Stone Bramble | *Rubus saxatilis* |  | x |  |
| Suffolk Lungwort | *Pulmonaria obscura* | x |  |  |
| Sweet Violet | *Viola odorata* |  | x |  |
| Sweet Woodruff | *Galium odoratum* | x |  |  |
| Thin Spiked Wood Sedge | *Carex strigosa* | x |  |  |
| Three-nerved Sandwort | *Moehringia trinervia* |  | x |  |
| Toothwort | *Lathraea squamaria* | x |  |  |
| Tunbridge Filmy-fern | *Hymenophyllum tunbridgense* | x |  |  |
| Tutsan | *Hypericum androsaemum* |  | x |  |
| Upland Enchanter's Nightshade | *Circaea x intermedia* |  | x |  |
| Violet Helleborine | *Epipactis purpurata* | x |  |  |
| Water Avens | *Geum rivale* |  | x |  |
|  |  | Woodland specialist +<br>slow colonizer | Woodland specialist<br>but known to colonize<br>secondary woodland | Generalist and/ or<br>swift colonizer |
| Water-purslane | *Lythrum portula* |  | x |  |
| White helleborine | *Cephalanthera damasonium* | x |  |  |
| Whitebeam | *Sorbus (microspecies)* |  |  | x |
| Wild Cherry | *Prunus avium* |  |  | x |
| Wild Daffodil | *Narcissus pseudonarcissus\** | x |  |  |
| Wild Service Tree | *Sorbus torminalis* | x |  |  |
| Wild Strawberry | *Fragaria vesca* |  | x |  |
| Wood Anemone | *Anemone nemorosa* |  | x |  |
| Wood Barley | *Hordelymus europaeus* | x |  |  |
| Wood Club-rush | *Scirpus sylvaticus* | x |  |  |
| Wood Crane's-bill | *Geranium sylvaticum* |  | x |  |
| Wood Fescue | *Festuca altissima* |  | x |  |
| Wood Forget-me-not | *Myosotis sylvatica* |  | x |  |
| Wood Horsetail | *Equisetum sylvaticum* |  |  | x |
| Wood Meadow-grass | *Poa nemoralis* |  | x |  |
| Wood Melick | *Melica uniflora* | x |  |  |
| Wood Millet | *Milium effusum* |  | x |  |
| Wood Small-reed | *Calamagrostis epigejos* |  | x |  |
| Wood Speedwell | *Veronica montana* |  | x |  |
| Wood Spurge | *Euphorbia amygdaloides* | x |  |  |
| Wood Stitchwort | *Stellaria nemorum* | x |  |  |
| Wood Vetch | *Vicia sylvatica* |  | x |  |
| Wood-sedge | *Carex sylvatica* |  |  | x |
| Wood-sorrel | *Oxalis acetosella* |  | x |  |
| Wych Elm | *Ulmus glabra* |  |  | x |
| Yellow Archangel | *Lamiastrum galeobdolon* | x |  |  |
| Yellow Bird's-nest | *Monotropa hypopitys* | x |  |  |
| Yellow Loosestrife | *Lysimachia vulgaris* |  |  | x |
| Yellow Pimpernel | *Lysimachia nemorum* |  | x |  |
| Yellow Star-of-Bethlehem | *Gagea lutea* |  | x |  |

**TABLE 4.** Deepcut Wood *WISDOM* Report (Woodland Indicator Strength, Distribution & Occurrence Matrix) SURVEYOR(S) AGB & JFL SURVEY DATE 26-28 April 2022 **Process** 1. Select the correct county AWI list for the survey area**Surrey** 2. Applicable species automatically selected below 3. Listed species differentiated into strong, moderate and weak indicators 4. Undertake botanical survey 5. Format *Distribution* columns to reflect compartment mapping & record occurrences accordingly, including totals in col. P 6. Assess indicator occurrences and report significance based on the following factors: a) Survey month: more indicators expected during M-J-J b) Size of woodland/ reporting area: more indicators expected in larger areas c) Topographic heterogeneity and presence of microclimates: more indicators expected with greater environmental variation Indicator strength has been assigned based on species’ known fidelity to ancient woodland and their frequency of occurrence (following a review of updated ancient woodland inventories).

| Scientific Name | Common Name | Indicator strength |  |  |  | Occurrence |
| --- | --- | --- | --- | --- | --- | --- |
|  |  | Strong | Moderate | Weak |  |  |
| * Caution required if specimen found within woodland edge: risk of garden escape or seed transport from outside woodland |  | Woodland specialist + slow colonizer | Woodland specialist but known to colonize secondary woodland | Generalist and/ or swift colonizer | S.M.W tally | 3.11.5 |
|  | List totals | 40 | 53 | 7 |  |  |
| Acer campestre* | Field Maple |  |  | x |  | R |
| Anemone nemorosa | Wood Anemone |  | x |  |  | R |
| Carex pendula* | Pendulous Sedge |  | x |  |  | O |
| Carex sylvatica | Wood-sedge |  |  | x |  | O |
| Carpinus betulus* | Hornbeam |  | x |  |  | R |
| Convallaria majalis* | Lily-of-the-Valley | x |  |  |  | R |
| Dryopteris affinis | Scaly Male-fern |  | x |  |  | R |
| Frangula alnus | Alder Buckthorn | x |  |  |  | R |
| Hyacinthoides non-scripta | Bluebell |  |  | x |  | R |
| Hypericum androsaemum | Tutsan |  | x |  |  | O |
| Ilex aquifolium | Holly |  |  | x |  | F |
| Lysimachia nemorum | Yellow Pimpernel |  | x |  |  | R |
| Polypodium vulgare (sensu | Polypody (all species) | x |  |  |  | R |
| Populus tremula* | Aspen |  | x |  |  | R |
| Potentilla sterilis | Barren Strawberry |  | x |  |  | R |
| Prunus avium | Wild Cherry |  |  | x |  | R |
| Ribes rubrum* | Red Currant |  | x |  |  | R |
| Rosa arvensis | Field-rose |  | x |  |  | R |
| Vaccinium myrtillus | Bilberry |  | x |  |  | O |

**TABLE 5.**
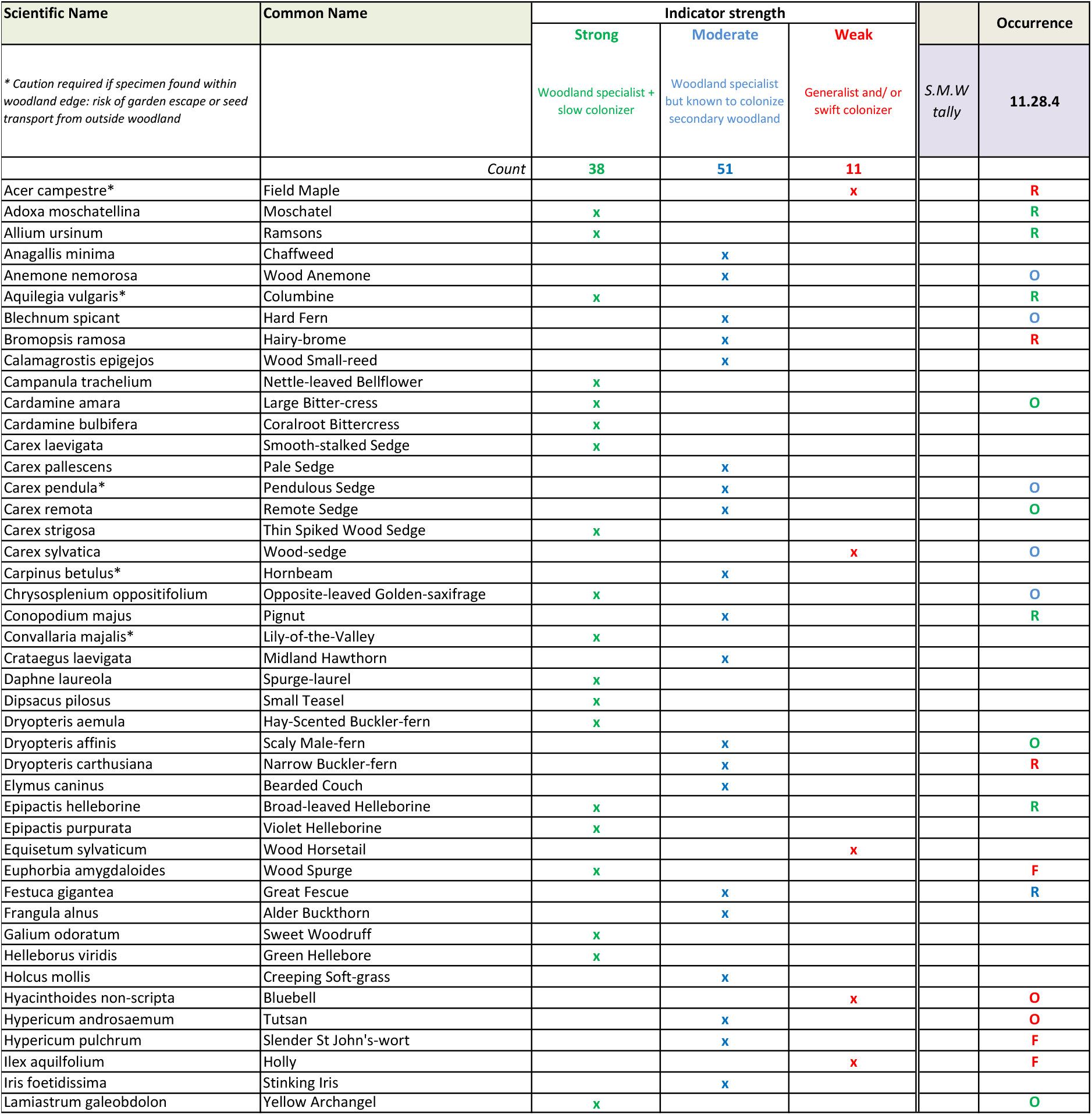
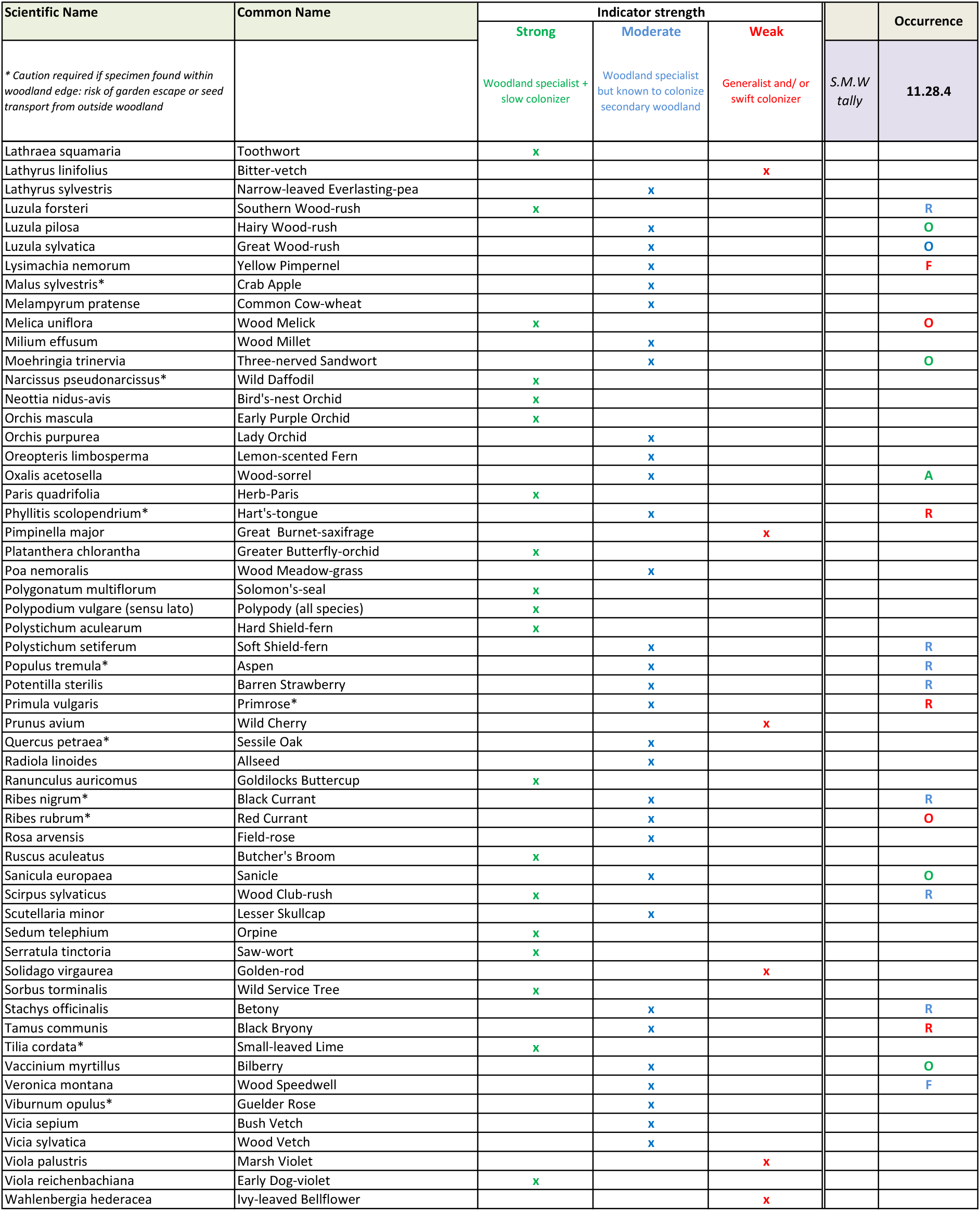
Verdley Wood *WISDOM* Report (Woodland Indicator Strength, Distribution & Occurrence Matrix) SURVEYOR(S) AGB & JFL DATE 27-30 June 2022 **Process** 1. Select the correct county or region AWI list for the survey area => 2. Applicable species automatically selected below 3. Listed species differentiated into strong, moderate and weak indicators 4. Undertake botanical survey 5. Format *Distribution* columns to reflect compartment mapping & record occurrences accordingly, including totals in col. P 6. Assess indicator occurrences and report significance based on the following factors: a) Survey month: more indicators expected during M-J-J b) Size of woodland/ reporting area: more indicators expected in larger areas c) Topographic heterogeneity and presence of microclimates: more indicators expected with greater environmental variation *Indicator strength has been assigned based on species’ known fidelity to ancient woodland and their frequency of occurrence (following a review of updated ancient woodland inventories)*.

| Scientific Name | Common Name | Indicator strength |  |  |  | Occurrence |
| --- | --- | --- | --- | --- | --- | --- |
|  |  | Strong | Moderate | Weak |  |  |
| * Caution required if specimen found within woodland edge: risk of garden escape or seed transport from outside woodland |  | Woodland specialist + slow colonizer | Woodland specialist but known to colonize secondary woodland | Generalist and/ or swift colonizer | S.M.W tally | 11.28.4 |
|  | Count | 38 | 51 | 11 |  |  |
| Acer campestre* | Field Maple |  |  | x |  | R |
| Adoxa moschatellina | Moschatel | x |  |  |  | R |
| Allium ursinum | Ramsons | x |  |  |  | R |
| Anagallis minima | Chaffweed |  | x |  |  |  |
| Anemone nemorosa | Wood Anemone |  | x |  |  | O |
| Aquilegia vulgaris* | Columbine | x |  |  |  | R |
| Blechnum spicant | Hard Fern |  | x |  |  | O |
| Bromopsis ramosa | Hairy-brome |  | x |  |  | R |
| Calamagrostis epigejos | Wood Small-reed |  | x |  |  |  |
| Campanula trachelium | Nettle-leaved Bellflower | x |  |  |  |  |
| Cardamine amara | Large Bitter-cress | x |  |  |  | O |
| Cardamine bulbifera | Coralroot Bittercress | x |  |  |  |  |
| Carex laevigata | Smooth-stalked Sedge | x |  |  |  |  |
| Carex pallescens | Pale Sedge |  | x |  |  |  |
| Carex pendula* | Pendulous Sedge |  | x |  |  | O |
| Carex remota | Remote Sedge |  | x |  |  | O |
| Carex strigosa | Thin Spiked Wood Sedge | x |  |  |  |  |
| Carex sylvatica | Wood-sedge |  |  | x |  | O |
| Carpinus betulus* | Hornbeam |  | x |  |  |  |
| Chrysosplenium oppositifolium | Opposite-leaved Golden-saxifrage | x |  |  |  | O |
| Conopodium majus | Pignut |  | x |  |  | R |
| Convallaria majalis* | Lily-of-the-Valley | x |  |  |  |  |
| Crataegus laevigata | Midland Hawthorn |  | x |  |  |  |
| Daphne laureola | Spurge-laurel | x |  |  |  |  |
| Dipsacus pilosus | Small Teasel | x |  |  |  |  |
| Dryopteris aemula | Hay-Scented Buckler-fern | x |  |  |  |  |
| Dryopteris affinis | Scaly Male-fern |  | x |  |  | O |
| Dryopteris carthusiana | Narrow Buckler-fern |  | x |  |  | R |
| Elymus caninus | Bearded Couch |  | x |  |  |  |
| Epipactis helleborine | Broad-leaved Helleborine | x |  |  |  | R |
| Epipactis purpurata | Violet Helleborine | x |  |  |  |  |
| Equisetum sylvaticum | Wood Horsetail |  |  | x |  |  |
| Euphorbia amygdaloides | Wood Spurge | x |  |  |  | F |
| Festuca gigantea | Great Fescue |  | x |  |  | R |
| Frangula alnus | Alder Buckthorn |  | x |  |  |  |
| Galium odoratum | Sweet Woodruff | x |  |  |  |  |
| Helleborus viridis | Green Hellebore | x |  |  |  |  |
| Holcus mollis | Creeping Soft-grass |  | x |  |  |  |
| Hyacinthoides non-scripta | Bluebell |  |  | x |  | O |
| Hypericum androsaemum | Tutsan |  | x |  |  | O |
| Hypericum pulchrum | Slender St John's-wort |  | x |  |  | F |
| Ilex aquilfolium | Holly |  |  | x |  | F |
| Iris foetidissima | Stinking Iris |  | x |  |  |  |
| Lamiastrum galeobdolon | Yellow Archangel | x |  |  |  | O |
|  |  | Strong | Moderate | Weak |  |  |
| * Caution required if specimen found within woodland edge: risk of garden escape or seed transport from outside woodland |  | Woodland specialist + slow colonizer | Woodland specialist but known to colonize secondary woodland | Generalist and/ or swift colonizer | S.M.W tally | 11.28.4 |
| Lathraea squamaria | Toothwort | x |  |  |  |  |
| Lathyrus linifolius | Bitter-vetch |  |  | x |  |  |
| Lathyrus sylvestris | Narrow-leaved Everlasting-pea |  | x |  |  |  |
| Luzula forsteri | Southern Wood-rush | x |  |  |  | R |
| Luzula pilosa | Hairy Wood-rush |  | x |  |  | O |
| Luzula sylvatica | Great Wood-rush |  | x |  |  | O |
| Lysimachia nemorum | Yellow Pimpernel |  | x |  |  | F |
| Malus sylvestris* | Crab Apple |  | x |  |  |  |
| Melampyrum pratense | Common Cow-wheat |  | x |  |  |  |
| Melica uniflora | Wood Melick | x |  |  |  | O |
| Milium effusum | Wood Millet |  | x |  |  |  |
| Moehringia trinervia | Three-nerved Sandwort |  | x |  |  | O |
| Narcissus pseudonarcissus* | Wild Daffodil | x |  |  |  |  |
| Neottia nidus-avis | Bird's-nest Orchid | x |  |  |  |  |
| Orchis mascula | Early Purple Orchid | x |  |  |  |  |
| Orchis purpurea | Lady Orchid |  | x |  |  |  |
| Oreopteris limbosperma | Lemon-scented Fern |  | x |  |  |  |
| Oxalis acetosella | Wood-sorrel |  | x |  |  | A |
| Paris quadrifolia | Herb-Paris | x |  |  |  |  |
| Phyllitis scolopendrium* | Hart's-tongue |  | x |  |  | R |
| Pimpinella major | Great Burnet-saxifrage |  |  | x |  |  |
| Platanthera chlorantha | Greater Butterfly-orchid | x |  |  |  |  |
| Poa nemoralis | Wood Meadow-grass |  | x |  |  |  |
| Polygonatum multiflorum | Solomon's-seal | x |  |  |  |  |
| Polypodium vulgare (sensu lato) | Polypody (all species) | x |  |  |  |  |
| Polystichum aculearum | Hard Shield-fern | x |  |  |  |  |
| Polystichum setiferum | Soft Shield-fern |  | x |  |  | R |
| Populus tremula* | Aspen |  | x |  |  | R |
| Potentilla sterilis | Barren Strawberry |  | x |  |  | R |
| Primula vulgaris | Primrose* |  | x |  |  | R |
| Prunus avium | Wild Cherry |  |  | x |  |  |
| Quercus petraea* | Sessile Oak |  | x |  |  |  |
| Radiola linoides | Allseed |  | x |  |  |  |
| Ranunculus auricomus | Goldilocks Buttercup | x |  |  |  |  |
| Ribes nigrum* | Black Currant |  | x |  |  | R |
| Ribes rubrum* | Red Currant |  | x |  |  | O |
| Rosa arvensis | Field-rose |  | x |  |  |  |
| Ruscus aculeatus | Butcher's Broom | x |  |  |  |  |
| Sanicula europaea | Sanicle |  | x |  |  | O |
| Scirpus sylvaticus | Wood Club-rush | x |  |  |  | R |
| Scutellaria minor | Lesser Skullcap |  | x |  |  |  |
| Sedum telephium | Orpine | x |  |  |  |  |
| Serratula tinctoria | Saw-wort | x |  |  |  |  |
| Solidago virgaurea | Golden-rod |  |  | x |  |  |
| Sorbus torminalis | Wild Service Tree | x |  |  |  |  |
| Stachys officinalis | Betony |  | x |  |  | R |
| Tamus communis | Black Bryony |  | x |  |  | R |
| Tilia cordata* | Small-leaved Lime | x |  |  |  |  |
| Vaccinium myrtillus | Bilberry |  | x |  |  | O |
| Veronica montana | Wood Speedwell |  | x |  |  | F |
| Viburnum opulus* | Guelder Rose |  | x |  |  |  |
| Vicia sepium | Bush Vetch |  | x |  |  |  |
| Vicia sylvatica | Wood Vetch |  | x |  |  |  |
| Viola palustris | Marsh Violet |  |  | x |  |  |
| Viola reichenbachiana | Early Dog-violet | x |  |  |  |  |
| Wahlenbergia hederacea | Ivy-leaved Bellflower |  |  | x |  |  |

Of the 19 AWIS found in Deepcut Wood, 14 (73.6%) were present either as single plants, or in single, small colonies; both of these occurrences were placed into the *Rare* DAFOR category.

Four species were placed into the *Occasional* DAFOR category, and one (Holly *Ilex aquifolium*) was noted to be *Frequent*. Holly is considered to be a weak indicator (poor fidelity to old woods with relatively rapid colony increase).

In contrast, of the 43 AWIS found in Verdley Wood, none were found as single plant or small colony occurrences, whilst one species was abundant (Wood sorrel *Oxalis acetosella*). Five species had frequent occurrence and 18 were occasional. The remaining 19 were placed into the *Rare* DAFOR category although, as noted, they were present in greater abundance than found at Deepcut Wood.

## Discussion

Whilst we acknowledge the small sample size of only two study woodlands, we consider that it can fairly be concluded that even a relatively high AWIS count (19 species in the case of Deepcut Wood) provides by itself no information regarding antiquity of woodland. Where, however, the AWIS are relatively numerous in both species and occurrence, this starts to provide useful information.

A very high AWIS count (>40 in this case) in confirmed ancient woodland is consistent with the indicator hypothesis, even though some of the species concerned were not present at high frequency.

Finally, our categorisation of evidential value as *Strong, Moderate* or *Weak*, renders it possible to consider a site’s AWIS assemblage in relative weighted terms. In this case, the known area of ancient woodland was found to have almost four times the *Strong* indicator presence than the known secondary woodland. We consider this to be more useful information than is provided by the presence of 19 AWIS in known secondary woodland.

## Conclusions

WISDOM improves the predictive power of AWIS lists and represents a step forward in the application of this technique. Having applied WISDOM in a comparative test of its efficacy for the investigation of landscape history, we are also able to draw the following broader conclusions in respect of ancient woodland indicators as a whole:

1. Our research confirms that quite young secondary woodland can host a substantial AWIS assemblage, even where the woodland in question is not connected in the landscape to ancient woodland. It follows that the presence of AWIS is not confirmatory of ancient woodland.
2. Of greater relevance than a simple species count are frequency of occurrence and relative evidential weight of the species concerned. In relation to these matters, WISDOM is of considerable assistance.
3. Where species count, occurrence and evidential weight align toward the positive, this is likely reflective of very long-established woodland. In other cases, the presence of AWIS needs careful interpretation. As a minimum, it should not be used to identify woodland as ancient. This is consistent with the position taken by Natural England (per reference 1).
4. Where woodland compartment mapping is undertaken, with individual WISDOM scores being recorded for each compartment, the distribution of indicators within the wider woodland (in addition to evidential weight and occurrence), can help identify remnants of ancient woodland surviving within a younger landscape.

The WISDOM spreadsheet can be downloaded free of charge from www.sylvan-consulting.com.

**Julian Forbes-Laird**

BA(Hons), Dip.GR.Stud, MICFor, MRSB, MRICS, MEWI, Dip.Arb(RFS)

**Alistair Baxter**

BA (Hons), MA (Oxon), MSc, CEcol, CEnv, MCIEEM

## Statement on the use of AI within this paper

The authors made use of the SciSummary journal paper summarizer tool (www.scisummary.com) to assist with the literature review contained in this paper. The tool was used chiefly as a test of its accuracy, insofar as the authors were already familiar with the contents of the papers concerned. After using the tool, the authors reviewed and edited the generated summary content as needed, and take full responsibility for the content included in this publication. It may assist others to know that we found the SciSummary tool to accurately reflect the contents of the papers tested. The authors have no relationship of any sort with the SciSummary team, and offer this observation merely to report its usefulness.

## Footnotes

1 https://www.exmoor-nationalpark.gov.uk/about-us/press-room/press-room/news-2022/bye-wood-bluebells-hint-at-ancient-wooded-past retrieved at 0900 on 15.06.23

2 *Handbook for Updating the Ancient Woodland Inventory for England*, NECR248, SANSUM, P. & BANNISTER, N.R. 2018

3 *Multi-Agency Geographic Information for the Countryside* https://magic.defra.gov.uk/

4 *Landscape Character Assessment of West Sussex*, West Sussex County Council 2003

